# Lung Tumor Microphysiological System with 3D Endothelium to Evaluate Modulators of T-Cell Infiltration

**DOI:** 10.1101/2022.08.03.502507

**Authors:** Katrina M. Wisdom, Johnny Suijker, Lenie van den Broek, BanuPriya Sridharan, Taraka Sai Pavan Grandhi, Aaron Cheng, Mahdi Lamb, Steven A. Titus, Andrew Gehman, Derek Poore, Niyant Shah, Shih-Hsun Cheng, Edward Kim, Sue Griffin, Jason Ekert

## Abstract

Lung cancer is a leading cause of death worldwide, with only a fraction of patients responding to immunotherapy. The correlation between increased T-cell infiltration and positive patient outcomes has motivated the search for therapeutics promoting T-cell infiltration. While transwell and spheroid platforms have been employed, these models lack flow and endothelial barriers, and cannot faithfully model T-cell adhesion, extravasation and migration through 3D tissue. Presented here is a 3D chemotaxis assay, in a lung tumor on chip model with 3D endothelium (LToC-Endo), to address this need. The described assay consists of a vascular tubule cultured under rocking flow, through which T-cells are added; a collagenous stromal barrier, through which T-cells infiltrate; and a chemoattractant/tumor compartment. Here, activated T-cells extravasate and infiltrate in response to gradients of rhCXCL11 and rhCXCL12. Adopting a T-cell activation protocol with a rest period enables proliferative burst prior to introducing T-cells into chips, increases T-cell expression of CXCR3 and CXCR4 receptors, and enhances assay sensitivity. In addition, incorporating this rest recovers endothelial activation in response to rhCXCL12. As a final control, we show that blocking ICAM-1 interferes with T-cell adhesion and chemotaxis. This microphysiological system, which mimics *in vivo* stromal and vascular barriers, can be used to evaluate potentiation of immune chemotaxis into tumors while probing for vascular responses to potential therapeutics. Finally, we propose a translational strategy by which this assay could be linked to preclinical and clinical models to support human dose prediction, personalized medicine, and the reduction, refinement, and replacement of animal models.

## Introduction

Although immunotherapy has shown great promise, immune cell infiltration into the tumor microenvironment of many indications and/or sub-indications remains challenging, leading to mixed clinical outcomes[1]–[3]. Patients with “inflamed” tumors, in which immune cells are inhibited but in close contact with tumor cells, typically respond better to cancer immunotherapy and experience better prognoses[4], [5]. By contrast, patients tend to experience poorer outcomes if their tumors are “immune excluded”, in which cytotoxic T-cells have accumulated in the tumor stroma but are not able to reach the tumor cells, or “immune desert”, in which cytotoxic T-cells are absent from both the tumor nest and stroma[4], [5]. Given that a high presence of cytotoxic T-cells in tumors is correlated with improved patient survival, there is a strong need to enhance T-cell chemotaxis into tumors and enhance the effectiveness of immunotherapies[1]–[3], [5], [6]. Despite the clear rationale to address this aspect of the cancer-immunity cycle, there are limited potential therapeutics available to address it [6].

While preclinical *in vivo* models have ushered in pivotal treatments in cancer immunotherapy (e.g., anti-CTLA-4 and anti-PD-(L)1), the limited translatability of preclinical models is a key challenge for the development of many immunotherapies[4]. Genetically engineered mouse models have evolved as the closest representation of human cancers, but differences in species-specific immunology and disease progression between mouse and human tumors hamper their clinical translatability[4], [7]. Furthermore, increasing global attention on ethical issues with animal research has bolstered support for initiatives to refine, reduce, and replace animal models[8].

*In vitro*, Transwell migration systems have been employed to investigate modulators of cell migration and chemotaxis. However, the effects of chemotactic triggers on migrating cells over long time windows remains challenging in these platforms due to gravity and gradient instability[9]–[11]. Furthermore, these platforms are unable to recapitulate some aspects of the tumor microenvironment. Transwell membranes with rigid pores are unable to model dynamic cell extravasation through living, responsive vasculature or 3D cell migration through viscoelastic and mechanically plastic pores of extracellular matrix[12]. Furthermore, as chemotaxis takes place along the z-axis in these assays, large confocal z-stacks, which may be time and data intensive to acquire and process, may be necessary to obtain single cell-resolution migration information. Alternatively, 3D spheroids are valuable for modeling T-cell infiltration into tumor nests [13]–[15]. However, optical clearing is necessary in order to image inside spheroids beyond 200 µm, which can only be done as an endpoint analysis. Furthermore, spheroid assays do not always model the extracellular matrix of solid tumors, even though dense stromal matrix is known to physically prevent infiltration in human lung tumors[16]. Additionally, growing evidence suggests that T-cells exhibit distinct kinds of motility dependent on both their activation state and features of their microenvironment[17]. For these reasons, infiltration studies with spheroids alone may not be sufficient to model the stromal constituents contributing to antitumor immunity in “immune excluded” and “immune desert” tumors.

While transwell and spheroid models can be informative and high throughput, they also lack a living endothelial barrier and vascular flow. For this reason, these platforms cannot be used to model extravasation, an early stage of T-cell chemotaxis into tumors. There is a need for an integrated complex *in vitro* model to investigate multiple stages of T-cell chemotaxis, including T-cell adhesion, extravasation, and infiltration through a 3D stromal barrier, to evaluate therapeutics that could enable T-cells to overcome these barriers and directly contact tumor cells, thereby enhancing the effectiveness of immunotherapies. For maximum utility in drug discovery and development, it should be phenotypic-screening amenable, offering single cell resolution readouts without being time and data intensive to image or analyze.

Recent developments in organ-on-chip technologies have been encouraging, but many of these early models are low throughput, made of polydimethylsiloxane (PDMS) (a hydrophobic material known to nonspecifically adsorb proteins), and contain artificial membranes[18]. The MIMETAS 3-lane Organoplate® is a platform containing 40 chips per plate, no PDMS, and phase guide technology, which enables membrane-free material separation. Recently, this platform was used to investigate monocyte-to-endothelium adhesion, neutrophilic infiltration, and 3D T-Cell chemotaxis in a melanoma model[18]. Building upon these models, we established a lung tumor on chip model with 3D endothelium (“LToC-Endo”) to investigate immune cell chemotaxis in response to chemokines and antibody treatments. Here we show that activated T-cells in the LToC-Endo model adhere, extravasate, and infiltrate in response to gradients of rhCXCL11 and rhCXCL12 (referred to throughout the manuscript as “CXCL11” and “CXCL12”), and that simultaneously, the living endothelial barrier responds to CXCL12 by sprouting. Using this assay, we show functional differences between T-cells activated using different approaches, and can inhibit infiltration by perturbing canonical endothelial receptor-T-cell receptor interactions.

## Results and Discussion

### Tumor barrier limits chemokine diffusion throughout tumor chips with 3D endothelium

We developed a lung tumor on chip model with 3D endothelium (“LToC-Endo”) in the MIMETAS 3-lane Organoplate® using three human cell types: pooled donor human umbilical vein endothelial cells (HUVEC), HCC0827 non-small cell lung carcinoma cell line, and primary T-cells. First, we established a collagen-1 extracellular matrix barrier. Then, we seeded endothelial cells in the top channel of the Organoplate® against this barrier on day −2, and cultured the chips under rocking flow (Fig 1a). The following day (day −1), we seeded tumor cells and by day 0, we observed that both the endothelial cells and tumor cells formed tubules in the top and bottom lanes, respectively (Fig 1b).

**Figure 1:**
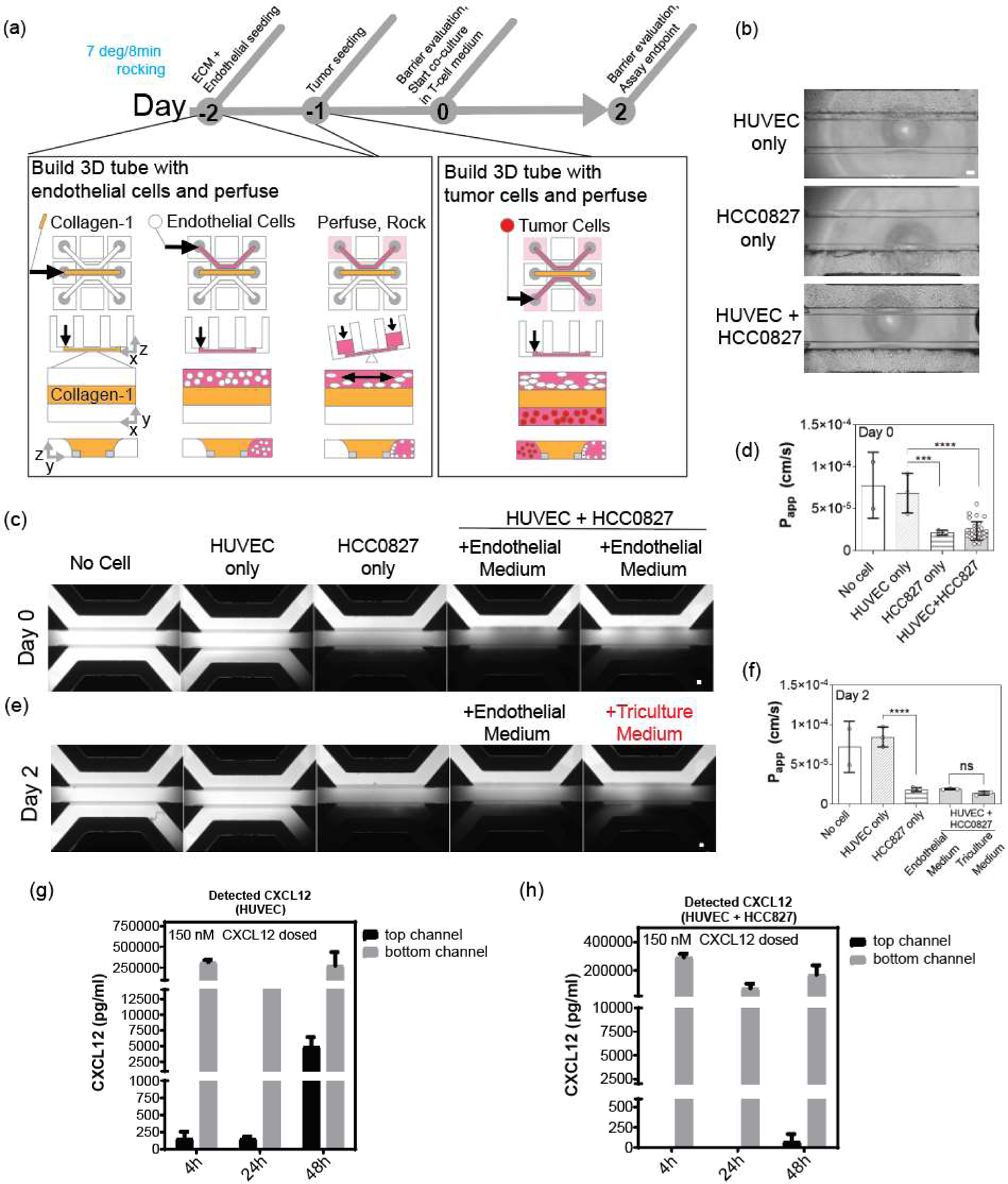
T-cell chemotaxis assay development, medium evaluation, and barrier analysis in the LToC-Endo model. (a) Experimental setup and timeline of platform seeding with Collagen-1, endothelial cells, and tumor cells. (b) Representative images of the platform seeded in monoculture and coculture configurations, with endothelial medium and triculture assay medium, on day 0. The transition from endothelial medium to triculture assay medium occurs on day 0 to mimic T-cell seeding at that time point. Refer to Fig S1 for information on the different assay media formulations considered. (c) and (e), Barrier integrity assay fluorescence images, where white shows FITC dextran presence. (d) and (f), permeability coefficient measurements for different configurations of the assay, for day 0 and day 2, respectively. Data show measurements per chip for n=2 or n>2 chips per condition for those used for statistical testing, bars indicate means, and error bars indicate standard deviations. Data was square root transformed prior to statistical testing to account for unequal variances. Outcomes are indicated for statistical tests comparing barrier diffusivity among the conditions tested (One-way ANOVA, ***p< 0.001, ****p<0.0001). (g) and (h), ELISA data of CXCL12 concentration in the bottom and top channels 48h after 150 nM CXCL12 is first introduced into the bottom channels of the chips. Data show measurements per chip for n = 2 chips per condition. The bars represent means and error bars indicate SD, and error bars indicate standard deviations. The scale bars in (b), (c), and (e) are 100 µm.

To evaluate the diffusivity of both the endothelial and tumor barriers in the LToC-Endo, we performed two different assays. First, using an imaging-based barrier integrity assay, we added 20 kDa fluorescent dextran (approximately the size of chemotactic chemokines) on day 0 into the top endothelial channel, and observed dextran flow through the chips over time with fluorescence microscopy (Fig 1c). By comparing permeability coefficients throughout different chip configurations, we noticed that the 3D endothelial tube readily allowed diffusion, and was comparable to no-cell chip controls (Fig 1c-f). The tumor tubule formed a more diffusion-limiting barrier than the endothelial cells, explaining why the combination of barriers is also significantly more diffusion-limiting than the endothelial barrier alone (Fig 1c-f). This finding was corroborated by an ELISA-based permeability assay, in which CXCL12 chemokine was added into the bottom channel, such that it flowed into the tumor tubule or empty channel and then diffused upward through the chip (Fig 1g,h). Media sampling over time revealed that in the no-tumor version of the assay, chemokine was detectable in the top channel as early as 4 hours, and increased markedly by the 48h time point, with a gradient remaining by this time. By contrast, with a tumor tubule, we could not detect any chemokine in the top channel after 48 hours, at which point the chemokine concentration in the top channel was comparable to, or less than, that of the no-tumor assay after only 4 hours. Altogether, these diffusion studies support that this assay models leaky tumor vasculature near a diffusion-limiting NSCLC tumor.

To make the LToC-Endo amenable to T-cell addition, we explored the impact of an assay medium switch on day 0 and then evaluated the corresponding platform permeability in addition to endothelial cell number and phenotype (Fig 1b,e, and f; Fig S1). We selected AIM V medium, supplemented with 5 ng/mL recombinant human VEGF (165 isoform) and bFGF, based on its ability to promote markers of endothelial tube stability without appreciably changing barrier diffusion properties.

### Activated, but not naïve, T-cells infiltrate in response to chemokine gradients in the tumor chips with 3D endothelium

Next, we used the LToC-Endo model to study the effect of activation status, T-Cell seeding density, tumor barrier presence, and chemoattractant type on T-cell chemotaxis. On day 0, we seeded either naïve or activated primary human T-cells into the endothelial channel of the tumor chip, along with recombinant CXCL11 or CXCL12 in the bottom tumor channels (Fig 2a). While naïve T-Cells did not appear to infiltrate into the ECM compartment, activated T-Cells infiltrated the ECM compartment in a seeding density-dependent manner by the day 2 time point after T-cell seeding, in response to both chemokines (Fig 2b-d, Fig. S2). We observed significant differences between chemokine and vehicle control chips (Fig 2c,d). These data are consistent with ELISA data previously shown, illustrating limited chemokine diffusion to the top channel until a 48-hour time point when a tumor barrier is present (Fig 1h).

**Figure 2:**
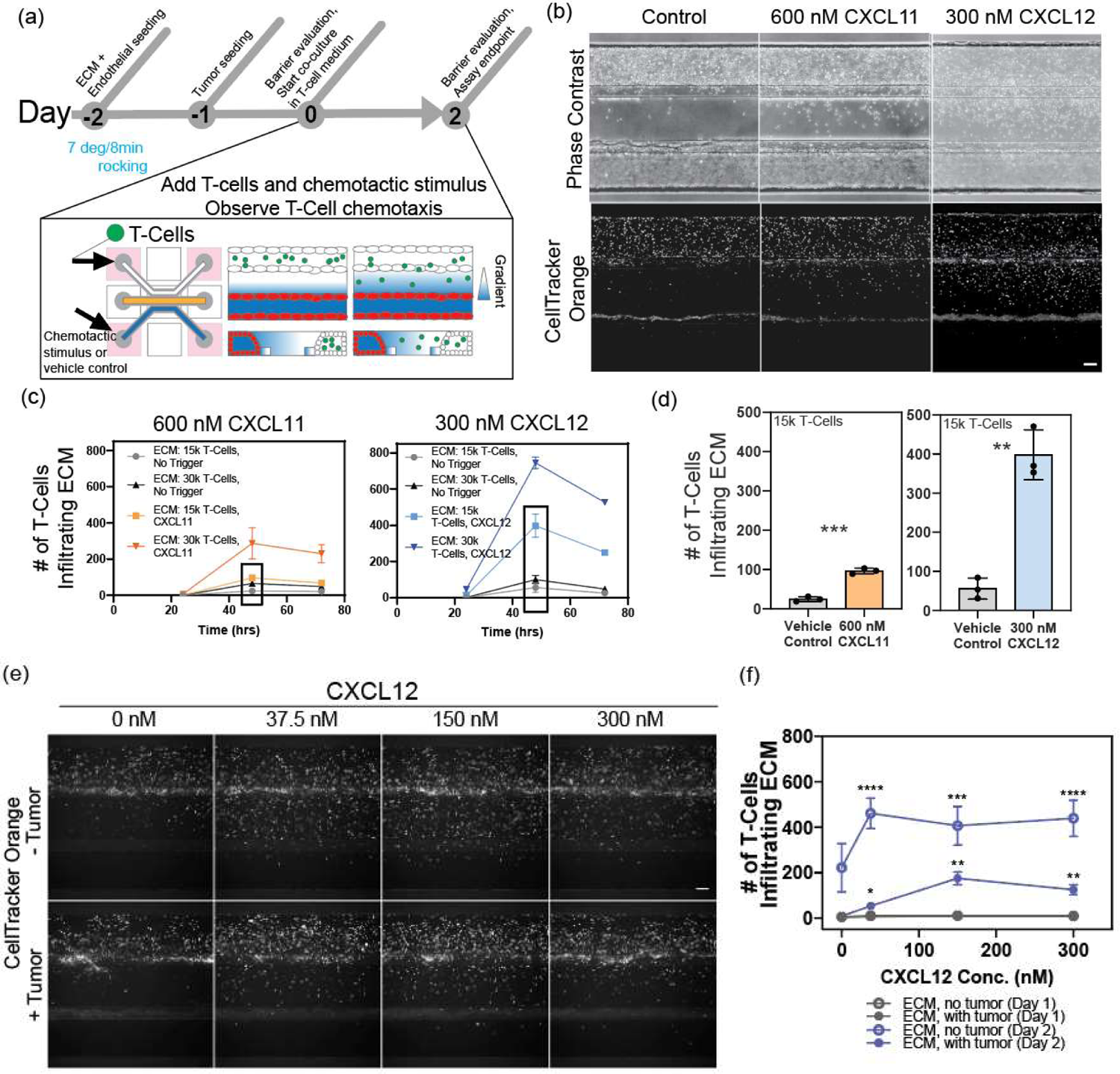
T-cell seeding density, chemokine type and dose, and tumor barrier presence regulate T-cell infiltration in the LToC-Endo model. (a) Experimental setup and timeline with platform seeding of extracellular matrix, endothelial cells, tumor cells, and T-cells. (b) Representative phase contrast and fluorescence images of T-cell infiltration into the collagen barrier of the tumor on chip, in response to chemokines CXCL11, CXCL12, vehicle controls. Images depict data using a T-cell seeding density of 15,000 cells/chip. (c) Number of infiltrated T-cells, by seeding density and over time, in CXCL11 and CXCL12 and respective plate controls, with data points indicating means of n =3 chips and error bars indicating SD. 48-hour time point data is highlighted in (d), with bars indicating means of n = 3 chips per condition and error bars indicating SD. Results shown for Welch’s t-tests to accommodate unequal variances (** p < 0.01, *** p < 0.001). (e) Representative fluorescent images of T-cell infiltration, for chips by dose of CXCL12 chemokine, with and without tumor barriers, at the day 2 time point. (f) Number of infiltrated T-cells by CXCL12 chemokine dose, with and without tumor barriers, at Day 1 and Day 2 time points. Day 1 data for tumor and no tumor conditions are overlapping. Markers indicate means of n ≥ 6 chips per condition, and error bars indicate SEM. Significant differences between CXCL12 dosages and respective vehicle controls are shown (Brown-Forsythe and Welch ANOVA tests, corrected for multiple comparisons, *p< 0.05, ** p<0.01, *** p<0.001, **** p < 0.0001). Central channel scale bars in (b) and (e) are 100 µm.

We then evaluated the role of chemoattractant dose and tumor presence on T-cell adhesion and infiltration. T-cell adhesion to the 3D endothelial tube did not increase with CXCL12 concentration, although it did increase with time at all doses tested (Fig S3a). However, all doses of CXCL12 tested, regardless of tumor presence, led to significant differences in T-cell infiltration compared to control chips, with the no-tumor version of the assay leading to greater overall T-cell infiltration (Fig 2e,f). We especially noticed in no-tumor conditions an elevated baseline level of infiltration even in the absence of chemokine, compared to the with-tumor assay (Fig 2e,f). One potential disease-relevant explanation for this could be soluble inhibitory factors secreted by the tumor cells. However, another explanation could be asymmetry of media consumption in the no-tumor version of the assay (i.e. endothelial cells and T-cells only, and present in the top channel, with a cell-free bottom channel), which may lead to a nutrient gradient that initiates nonspecific T-Cell infiltration even in the absence of recombinant chemokine. What is more, ELISA-based diffusion studies repeated with T-cells support that the no-tumor version of the assay is more permissive to chemokine diffusion at all doses of chemokine tested (Fig S3b,c). Thus, more effective chemokine diffusion may also explain why T-cell response saturates at lower doses in the notumor assay (37.5 nM CXCL12) compared to the with-tumor version of the assay (150 nM CXCL12) (Fig 2f).

### Presence and activation of T-cells influence endothelial activation in response to CXCL12

In the LToC-Endo, we observed notable differences in endothelial tube response to CXCL12 depending on the presence of activated T-cells. While CXCL12 drives migration or angiogenic sprouting with naïve T-cells (Fig 3a) or when T-cells were absent (Fig 3b,c), we did not observe pervasive endothelial cell activation when introducing activated T-cells (Fig 3a). CXCL12 is a known driver of T-cell chemotaxis, but it is also a crucial regulator of angiogenesis. It acts by increasing VEGF-A production in endothelial cells, which then upregulates their CXCR4 expression, enhances responsiveness to CXCL12, and contributes to an amplifying angiogenic signaling loop[19]–[21]. CXCL12 also promotes angiogenesis through Akt activation via atypical CXCR7 receptors, which are overexpressed only in stressed endothelial cells[22]. Under proangiogenic signaling, the endothelium responds by increasing endothelial wall permeability, destabilizing the vessel wall, and increasing expression of leukocyte adhesion receptors, in addition to increasing endothelial cell proliferation and migration[21], [23]–[25]. It is possible that these CXCL12-mediated endothelial events indirectly contribute to the observed window in T-cell infiltration (Fig 2b-f), in addition to the direct effect of CXCL12 driving T-cell chemotaxis. By contrast, CXCL11 is an angiostatic chemokine, known to counterbalance the vascular changes described above[21]. Therefore, as expected we do not see angiogenic sprouting in response to this chemokine in the assay. (Fig 3).

**Figure 3:**
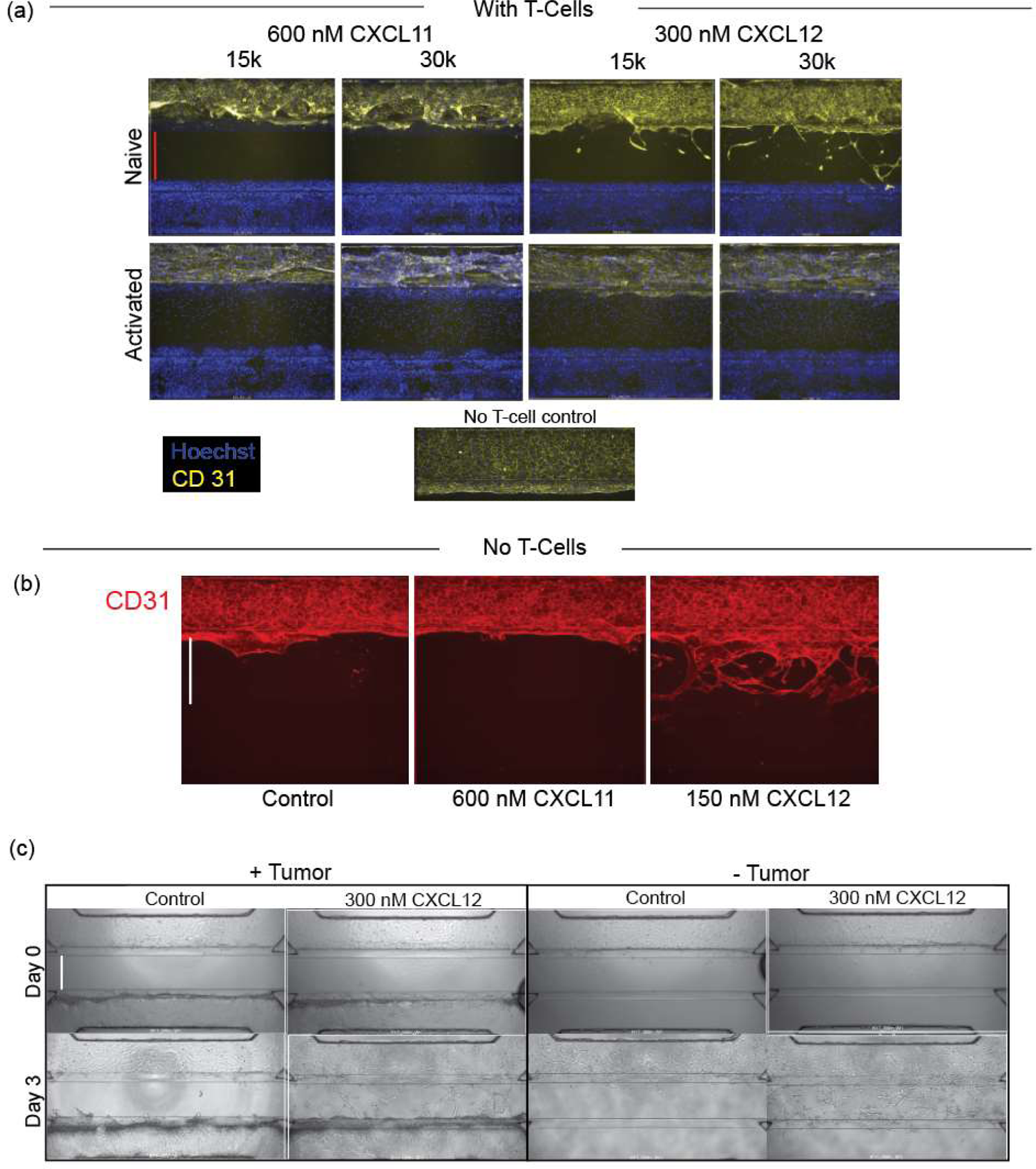
Migration and sprouting of 3D endothelium in response to rhCXCL12 in the LToC-Endo model. (a), Hoechst and CD31 staining of the indicated conditions, for naïve and activated T-cells, low and high T-cell seeding density, and CXCL11 and CXCL12 chemokines, on day 3. With no T-cells in the chips, (b) CD31 staining depicting 3D endothelium response to control, CXCL12, or CXCL11 conditions after 3 days in culture and (c) brightfield images showing endothelial response to CXCL12 or media control, with and without tumor cells, after 3 days in culture. In (a) through (c), middle channel width is 350 μm as indicated by the vertical bars.

The reduction in migration and angiogenic sprouting responsiveness to CXCL12 suggests that the 3D endothelium in the LToC-Endo may be under stress with the addition of activated T-cells. Images of the 3D endothelium, 3 days after activated T-cell addition, show large holes that are suggestive of endothelial stress (Fig 3a). Abundant T-cell proliferation is suspected to play a role, as both in-chip and off-chip T-cells exhibit an expected, post-activation proliferative burst (Fig S4), leading to a higher effective number of T-cells than initially seeded. As a consequence, the rapidly proliferating T-cells may not only be contributing to endothelial stress at long time points, but also diluting live cell dye, all of which may contribute to the plateau or decline in infiltrated T-cells after 2 days, which was seen both here (Fig 2c) and in a previous study [18].

### Alternate T-cell activation protocol impacts T-cell phenotype and enhances functional response

We hypothesized that the introduction of a rest period, mimicking the time lag between T-cell activation and homing to a tumor site *in vivo*[6], would allow us to overcome the proliferative burst prior to seeding activated T-cells. Additionally, we switched to a live nuclear dye which we expected would be stable over longer culture periods. We performed the assay side-by-side with activated or activated-rested T-cells. As expected, the activated-rested T-cells, which undergo proliferative burst during the 2-day rest, increase in concentration by 3-4x prior to seeding, compared to the activated-only T-cells (Fig S5a). Surprisingly, in spite of the lack of in-chip proliferative burst, activated-rested T-cells adhered to the endothelium and infiltrated in greater numbers in response to CXCL11 (∼4x more on average) and CXCL12 (∼2x more on average), compared to activated-only T-cells (Fig 4a-d). To better understand these changes, we profiled T-cells prepared using both approaches for expression of CXCR3 and CXCR4, the cognate receptors for CXCL11 and CXCL12. We saw that the introduction of a rest period following a T-cell activation enhanced CXCR3 and CXCR4 expression in all T-cell subsets compared to activation-only T-cells, increasing the overall proportion of double positive (CXCR3+CXCR4+) T-cells from ∼30-50% to ∼85% (Fig S5b, Table 1). Furthermore, we observed that the rest period led to more central and effector memory T-cell phenotypes, indicating a more durably activated state [26] (Fig S5c, Table 1). Our incorporation of an additional control in these studies allowed us to attribute changes in T-cell phenotype to differences in activation regimen, rather than culture medium, as this was also changed (Fig S5b,c; Table 1). Altogether, these data suggest that implementing a T-cell culture protocol with activation followed by a rest period enables the introduction of T-cells that are more sensitive and responsive into the tumor-on-chip assay.

**Figure 4:**
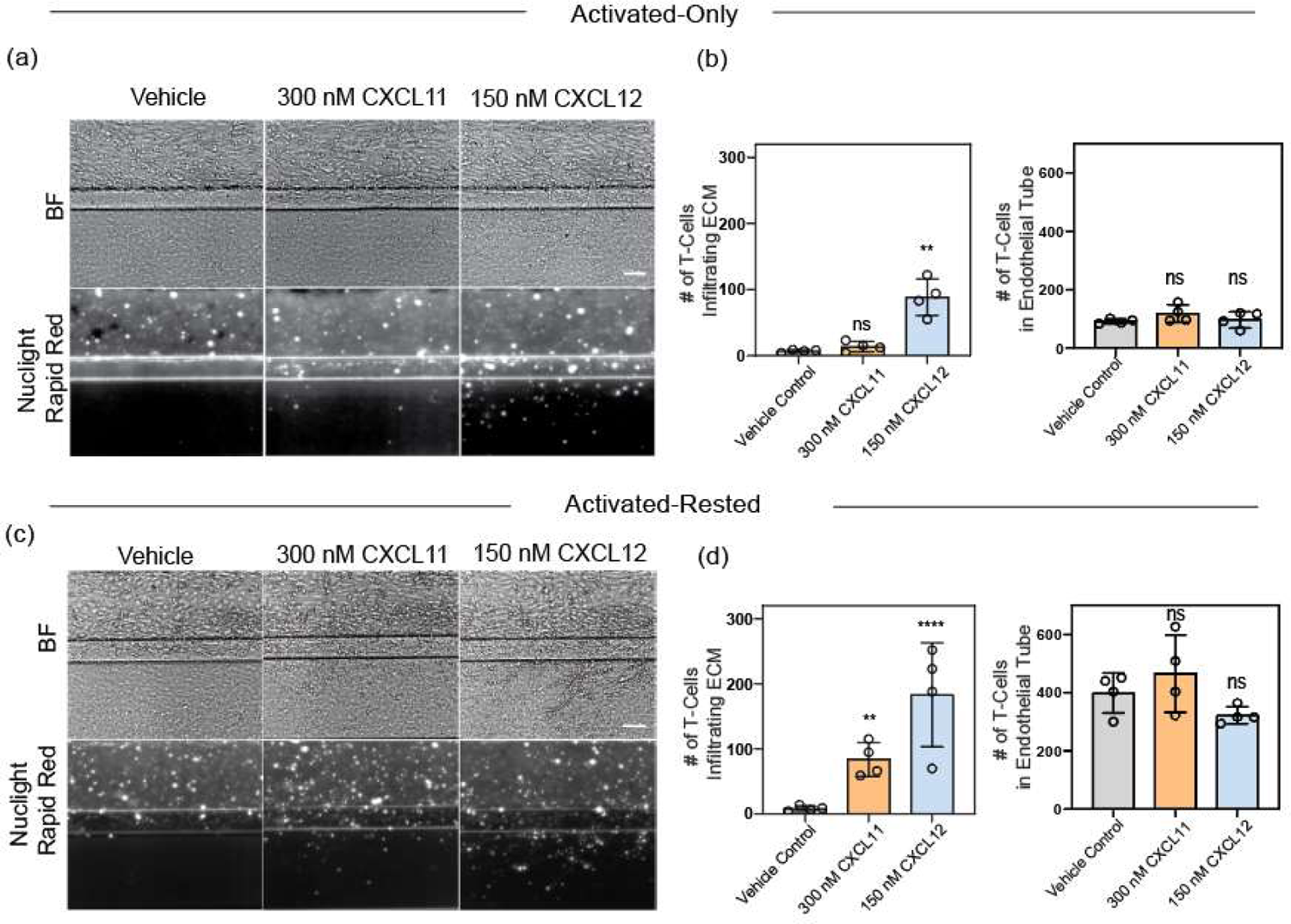
Activated-rested T-cell protocol enhances T-cell adhesion and chemotaxis, and restores CXCL12-driven endothelial activation, in the LToC-Endo. Representative brightfield and fluorescent images of T-cells (15k per chip) within the endothelial tubule and infiltration into the ECM compartment in response to the chemokine and dose indicated, at Day 2 time point, for (a) Activated-Only T-cells (AIMV) and (c) Activated-Rested T-cells (RPMI). Refer to supplement for flow cytometry controls that include additional Activated-Only (RPMI) condition. Scale bars in (a) and (c) are 100 µm. In (b) and (d), quantifications of infiltrated T-cells and T-cells within the endothelial tube for both T-cell preparation protocols, respectively. Markers indicate mean T-cell numbers per chip (n = 4 per condition), bars indicate mean T-cell numbers per condition, and error bars indicate SD. Statistical testing was performed on square root transformed data to satisfy criteria of equal standard deviations. Significant differences between chemokines and respective vehicle alone controls are shown (One-way ANOVA corrected for multiple comparisons, *p< 0.05, *** p<0.0001).

With respect to endothelial tube responsiveness to CXCL12 chemokine, another key difference emerged when switching from an activated-only to activated-rested T-cell culture protocol. With the activated-only T-cells, a lack of response previously shown (Fig 3a) was reproduced (Fig 4a), whereas in activated-rested T-cell chips, we observed endothelial migration in response to CXCL12 by 48h (Fig 4c). This endothelial response to CXCL12 with activated-rested T-cells appears to match more closely the endothelial response to CXCL12 with the naïve and no-T-cell conditions (Fig 3). These data lead us to infer that adding rested T-cells minimizes the stress on 3D endothelial tubes caused by T-cell addition, and may preserve more physiologically relevant responsiveness of the 3D endothelium to angiogenic cues.

### Assay timeline extension is facilitated by alternate T-cell activation protocol

Given that an activated-rested T-cell protocol allowed us to circumvent proliferative burst in-chip, preserve 3D endothelium responsiveness to activation, and mitigate live cell dye dilution, we hypothesized that we could extend the assay timeline. Repeating the assay with activated-rested T-cells, we compared day 2 and day 5 time points. While the activated-only T-cell version of the assay results in a decline in T-cell infiltration after day 2 (Fig 2c) and complete endothelial dissolution by day 5, the activated-rested T-cell version of the assay shows higher levels of T-cell adhesion and infiltration and more intact endothelium by day 5 (Fig 5a-d, Fig S6). While by day 2 we observe comparable levels of T-cell adhesion between control and chemokine conditions, by day 5 we observe significantly fewer T-cells adherent in the CXCL12 condition. This may reflect that although higher numbers of T-cells adhere to the endothelium over time in all conditions, a significant number have migrated due to extravasation in the CXCL12 condition (Fig 5d). Furthermore, we confirm that with this new T-cell activation strategy and extended timeline, T-cell adhesion and infiltration still scales with T-cell seeding density, as observed in prior studies (Fig 5e,f). Finally, T-cell infiltration remains greater when the tumor barrier is absent than when the tumor barrier is present, reflecting a trend previously observed (Fig 5c,e). Overall, these studies support that the adoption of an activated-rested T-cell culture protocol and a long-lasting, live nuclear dye enable assay timeline extension.

**Figure 5:**
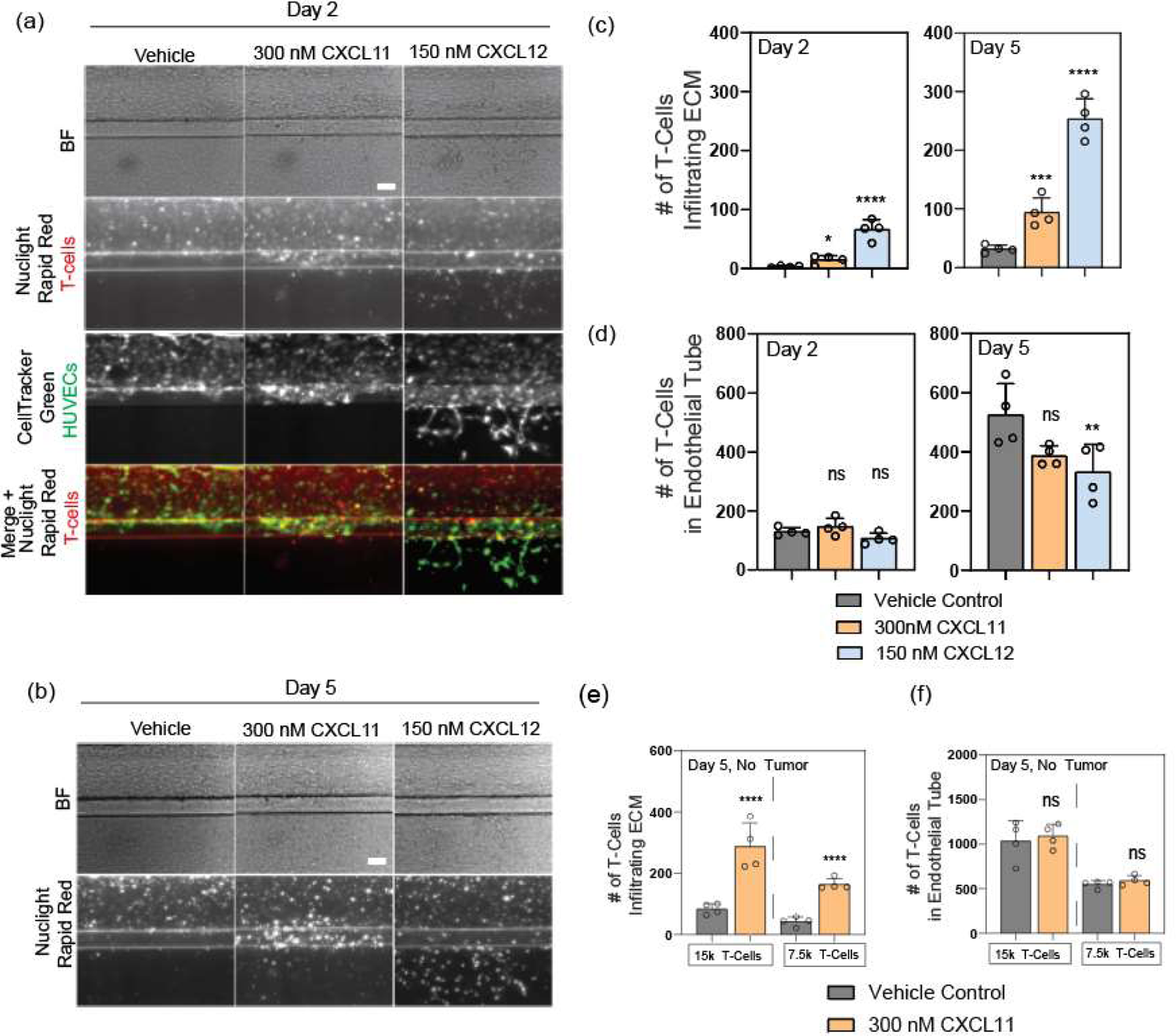
Activated-Rested T-cells enable an extended assay endpoint. (a) and (b), Representative brightfield and fluorescent images of T-cell infiltration in response to the indicated chemokines and doses, on Days 2 and 5, for assay with 15k T-cells seeded and with tumor barrier. Scale bars are 100 µm. (c), Quantifications of infiltrated T-cells and (d) T-cells within the endothelial tubes on Days 2 and 5, for the assay with tumor barrier. Day 5 quantifications from studies without tumor barriers, of (e) infiltrated T-cells and (f) T-cells within the endothelial tubes, for two different T-cell seeding densities. In (c) through (f), markers indicate T-cell numbers per chip (n = 4 per condition), bars indicate mean T-cell numbers per condition, and error bars indicate SD. In (c) and (e), statistical testing was performed on square root transformed data to satisfy criteria of equal standard deviations. Significant differences between chemokines and respective vehicle controls are shown (One-way ANOVA corrected for multiple comparisons, *p< 0.05, ** p<0.01, **** p<0.0001).

### T-cell chemotaxis in tumor on chip requires ICAM-1

Finally, we evaluated the ability of this tumor on chip microphysiological system to recapitulate mechanisms of T-extravasation. Tumor infiltration requires chemokine-induced polarization of T-cells and attachment to the endothelium through VCAM-1/ICAM integrin activity[27], [28]. Therefore, we repeated this assay using CXCL11 as the chemotactic stimulus, and added blocking antibodies against endothelial receptors VCAM-1 and ICAM-1 or isotype controls at the same time as adding chemotactic triggers.

By Day 5 of T-cell incorporation into the platform, we observe that ICAM-1 blocking antibody treatment significantly impacts the number of T-cells infiltrating into the extracellular matrix in response to CXCL11, while VCAM-1 blocking antibody does not (Fig 6a,b). The median infiltration distance of T-cells in chips with ICAM-1 blocking antibody is significantly reduced compared to those with isotype control (Fig 6c). These trends hold in with-tumor and without-tumor versions of the assay (Fig 6a-c). We do observe that the addition of IgG control antibody significantly impacts the number of T-cells adhering to the endothelium, even in the absence of chemokine (Fig 6d). In the presence of CXCL11, and in the with-tumor assay condition, we observe a significantly lower number of T-cells adhering to the endothelium using ICAM-1 blocking antibody. Altogether, these data suggest that blocking ICAM-1 is sufficient to reduce but not entirely block T-cell adherence, extravasation, and chemotaxis. These findings are consistent with T-cells using endothelial receptors other than ICAM-1 to adhere, extravasate, and infiltrate.

**Figure 6:**
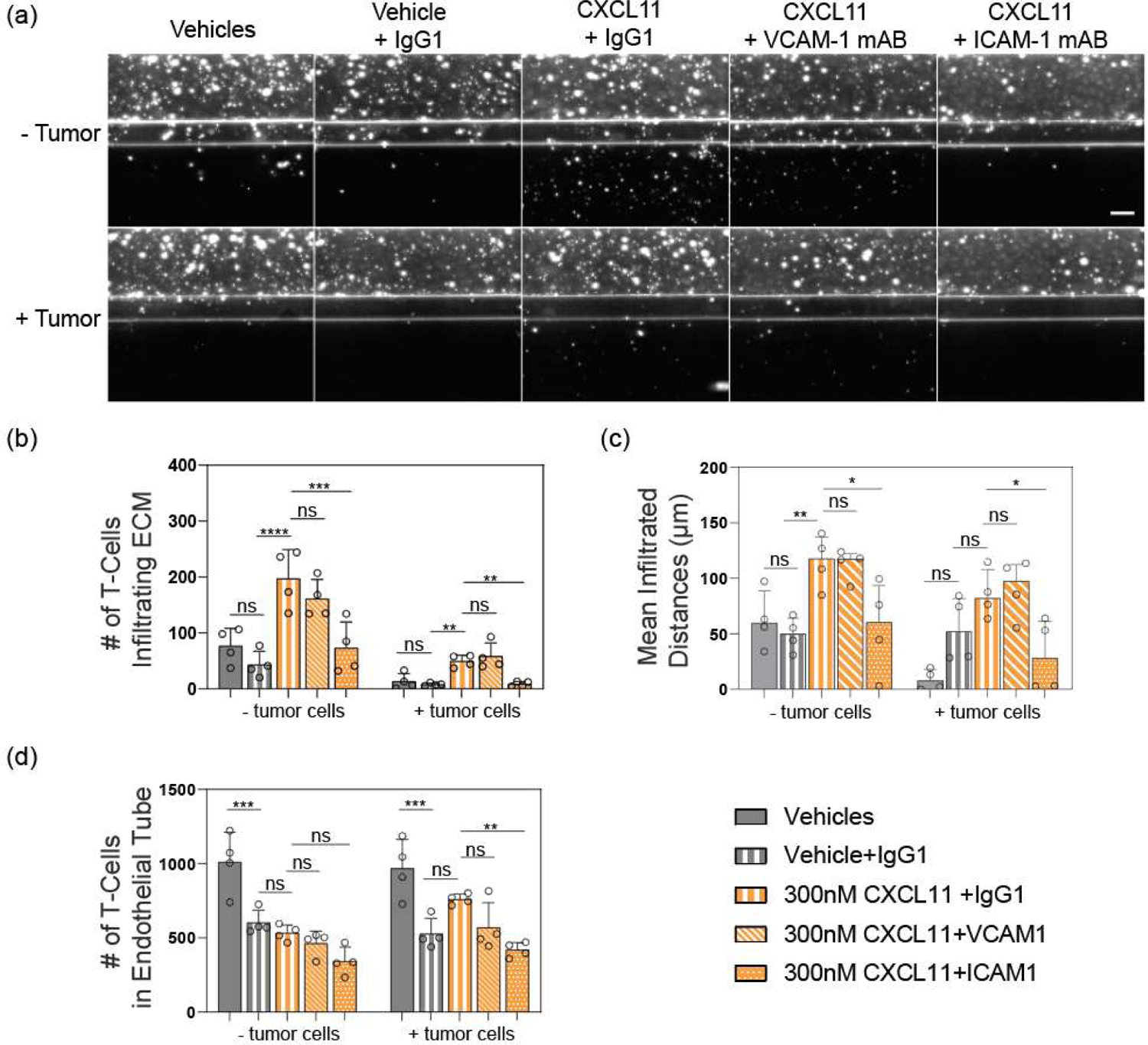
T-cell extravasation and chemotaxis in response to CXCL11 are dependent on ICAM-1 endothelial receptor in the tumor on chip platform. (a) Representative fluorescent images of T-cell infiltration in response to chemokine or vehicle control, with additional treatment as indicated with blocking antibody or IgG control. Images show Day 5 assay data both without and with tumor barrier. Scale bar is 100 µm. Per-chip day 5 quantifications of (b) mean infiltrated T-cell number, (c) median infiltrated distance, and (d) mean T-cell number within the endothelial tubes for all conditions tested. In (b-d), markers indicate T-cell numbers per chip (n = 4 per condition), bars indicate mean T-cell numbers, and error bars indicate SDs. Significant differences between chemokines and respective vehicle alone controls, with or without antibody treatments, are shown (One-way ANOVA corrected for multiple comparisons, *p< 0.05, ** p<0.01, *** p<0.001, **** p<0.0001). In (b), statistical testing was performed on square root transformed data to satisfy criteria of equal standard deviations.

It is unclear why VCAM-1 blocking did not result in decreased adhesion and chemotaxis. In preclinical animal models, VCAM-1 density and tumor perfusion are predictive of T-cell infiltration and treatment response to adoptively transferred and endogenous T-cells[28]. However, blocking VCAM-1 is only marginally effective at blocking T-cell adhesion to endothelial cells *in vivo*. By contrast, combined blocking of CD49d/integrin-α4 (a VCAM-1 binding partner), and CD18/integrin β2 (an ICAM binding partner) offers substantially improved blocking, with this cocktail shown to prevent T-cell mediated tumor rejection[28].

In vitro, the role of VCAM-1 and ICAM-1 in T-cell adhesion depends on both T-cell and endothelial cell activation[29]. While ICAM-1 is the main ligand utilized by CD4+ T-cells to adhere to IL-1-induced HUVECs, memory T-cells can leverage a variety of adhesion pathways to bind to HUVECs, including VCAM-1, ICAM-1, ELAM-1, and other ICAM ligands[29]. Upon T-cell phenotyping here, we noted a switch from activated-only to activated-rested T-cells leads to a shift toward central and effector memory phenotypes (Fig. S5c). More memory T-cells, utilizing a greater variety of adhesion pathways to achieve arrest on and extravasation through the endothelium, perhaps explains the enhanced T-cell adhesion and chemotaxis observed switching from the activated-only to activated-rested protocol (Fig 4b,d) as well as the partial blocking of T-cell infiltration using ICAM-1 blocking antibody (Fig. 6).

## Conclusion and Future Outlook

In conclusion, we developed a microfluidic lung tumor on chip assay with a 3D endothelium (LToC-Endo) perfused with rocking flow to evaluate modulators of T-cell extravasation and infiltration through 3D extracellular matrix in a non-small cell lung carcinoma (NSCLC) context. Due to the orientation of the platform, T-cell chemotaxis takes place across the x-y plane. This orientation readily facilitates snapshots of T-cell chemotaxis profiles across the stromal matrix, making the assay amenable to phenotypic screening and migration time point analysis. In alignment with a need for future work highlighted previously[18], we extended the assay timeline and improved the assay window by introducing a rest period after T-cell activation and selecting a long lasting, live nuclear dye. Similar to *in vivo*, activated T-cells in the LToC-Endo extravasate and infiltrate in response to chemotactic gradients, and the living endothelial barrier responds to pro-angiogenic cues through sprouting. We have also shown the dependence of T-cell infiltration on the presence of non-small cell lung carcinoma cells and on ICAM-1 endothelial receptors. While animal models typically recapitulate immune cold tumors, the LToC-Endo and described chemotaxis assay can also recapitulate features of immune-excluded tumors (i.e. angiogenesis, immune infiltration into stroma)[4]. Given differences in chemokines present and antigen-presenting functions of endothelial cells between human and animal models[4], this assay will be a valuable tool for probing humanized tumor-immune-endothelial multicellular interactions in NSCLC and other cancers. Additionally, this *in vitro* assay simultaneously offers the ability to observe compound efficacy (i.e. T-cell adhesion and infiltration) with safety (i.e. drug induced vascular injury, exacerbated angiogenesis in the tumor microenvironment (TME)), bringing safety information earlier into the discovery research pipeline.

Similar to what has been shown for an angiogenesis assay using this platform[30], the next step will be to evaluate the reproducibility and robustness in the LToC-Endo. Establishing a positive control with a clinically meaningful 2-5 fold window, yet without angiogenic side-effects, would be ideal based on prognostic differences between immune phenotypes in colorectal cancer tumors[31] and in alignment with robust assay design[32]. Further work is needed to validate the translatability of the assay by using standard of care molecules and comparing outcomes to clinical responses[33]. Moreover, there is a need to identify the T-cell subtypes that potential therapeutics successfully induce to infiltrate; in this case, enhancement of CD8+ cytotoxic T-cells is would be desirable. Future directions for the LToC-Endo involve incorporating the stromal cells that perpetuate immune suppression in the TME, such as cancer-associated fibroblasts, myeloid derived suppressor cells, and tumor-associated macrophages[34]–[36]. It will also be important to evaluate how other immune cell types (i.e. T-regulatory cells, natural killer cells, and B-cells[3]) infiltrate into the TME in response to chemotactic cues and compounds, and to include a tumor cell killing component into the assay. Altogether, we anticipate that the LToC-Endo complex in vitro model will serve as a valuable tool to study multicellular and cell-extracellular matrix mechanisms of immune suppression, screen for drug candidates that target these processes to improve patient responses to immunotherapies.

For this complex in vitro model to support the refinement, reduction, and replacement of animal immuno-oncology models, whether classical syngeneic (i.e. MC38, 4T1) or ‘humanized’ mouse tumor models[37], a translational strategy is needed. We propose that noninvasive imaging techniques serve as a translational link to align imaging-based pharmacodynamic (PD) timepoint readouts between complex in vitro and in vivo models of immune infiltration. Noninvasive imaging techniques can detect and monitor anatomical, functional, metabolic, or molecular-level changes within the body of animals with minimal pain, distress, or premature termination[38], and can do so in a temporal and spatial manner. For example, infiltration of specific T-cell populations (e.g. CD8+) can be tracked into specific organs, tumors, or tumor-draining lymph nodes over time within a single animal[39]. In this way, noninvasive imaging can enable comprehensive, longitudinal immune response datasets to be derived from fewer animals, thereby increasing the statistical power of the data gathered by reducing experimental variation[40]–[43].This is in contrast to traditional methods requiring animals to be sacrificed at given time points, i.e. using histology and flow cytometry[37]. Instead of relying exclusively on these informative yet endpoint-requirement techniques, which now include scRNAseq[41], they could instead be employed as-needed to verify or supplement noninvasive longitudinal imaging. Ideally, these noninvasive *in vivo* approaches would translate to evolving clinical imaging techniques, which are expected to gather similar longitudinal immune infiltration data, monitor therapeutic response in individual patients, and enable precision oncologic medicine[39]–[41].

With this translational strategy in mind, an imaging-based, humanized, immune infiltration complex in vitro model such as the LToC-Endo would be well suited to establish an in vitro/in vivo correlation. Longitudinal, imaging-based T-cell infiltration datasets, gathered per-chip, per-animal, and per patient, could then be used to calibrate silico models, enable better in vivo response prediction, refine the selection of candidates to progress into animal studies, and ultimately provide better medicines to patients.

## Data Availability Statement

The datasets generated during and analyzed during the current study are available from the corresponding author upon reasonable request.

## Author Contributions

K.M.W., B.S., J.E., A.C., E.K., D.P., N.S., J.S., and L.v.d.B. conceived of ideas and planned experiments. K.M.W., J.S., and L.v.d.B. led and performed key experiments. J.S. and K.M.W. analyzed experimental data. D.P. and N.S. prepared T-cells and performed T-cell receptor expression and phenotyping studies in GSK experiments. B.S., T.S.P.G, A.C., S.G., E.K., D.P.,and N.S. provided input and feedback on experimental studies. S.T. designed high content imaging protocols and data acquisition workflows for GSK experiments. M.L. designed GSK automated analysis pipeline for experimental data. A.G. provided recommendations for data analysis and statistical testing. K.M.W. prepared and wrote the manuscript, and S.G., T.S.P.G., J.E., N.S., and S.-H.C. contributed to the final manuscript.

## Acknowledgements

We gratefully acknowledge John Lowman for his facilitation of the collaborative research agreement. Thank you to Kimberly Nadwodny and Ed Gimmi for their data integrity and patents review, respectively. We also acknowledge Maggie Connelly for isolating T-cells for these studies. Thank you to Jeremy Waight for providing helpful feedback on the presentation of this work and the manuscript. This work was funded by GSK.

## Ethics Statement

The human biological samples were sourced ethically, and their research use was in accord with the terms of the informed consents under an IRB/EC approved protocol.

## Materials and Methods

### Cell Culture and Media

All human biological samples were sourced ethically, and their research use was in accord with the terms of the informed consents under an IRB/EC approved protocol. Human Umbilical Vein Endothelial Cells (HUVECs) (Lonza, pooled donor) were cultured in complete human endothelial medium (Cell Biologics), expanded, and bio-banked in aliquots. HUVECs in all studies were used at or before passage 5. HCC0827 cells (University of Texas Southwestern) were cultured in RPMI (Gibco) + 5% FBS (Gibco). Primary human T-Cells (Peripheral Blood, Cryopreserved, CD3+ Pan T Cells, Negatively Selected CD 3+, AllCells) were thawed in one of the following media solutions, as indicated in the studies described: either AIM V medium (Gibco) containing 20 IU/mL of IL-2 (Miltenyi) or RPMI + 10% FBS. For activated T-Cells, 1:500 TransAct (Miltenyi) was added to the medium. Activated-only T-cells were cultured for 48 hours (either with or without 1:500 TransAct) prior to use in assay. Activated-rested T-Cells were cultured for 72 hours in 1:500 TransAct, followed by a 48-hour rest period, during which time the medium was washed out via centrifugation and replaced with RPMI + 10% FBS.

### T-Cell Isolation

T-cells were obtained directly from AllCells and shipped to the MIMETAS research facility, or they were isolated from AllCells leukopaks internally at GSK. For this, T-cells were isolated from full fresh leukopaks (AllCells). Leukopaks were received and stored at 4C overnight (approx. 16h). First, Peripheral Blood Mononuclear cells (PBMCs) were isolated using a Custom PBMC Isolation Kit (Miltenyi), using magnetic beads to isolate out erythrocytes and granulocytes on magnetically charged cell selection columns while eluting PBMCs. T-cells were then isolated from the PBMCs using a standard Pan T Isolation kit (Miltenyi) using manufacturer protocols. T-cells were cryopreserved in CS10 (BioLife Solutions, 210102) in a rate-controlled freezer over the course of one hour, and transferred to LN2 storage.

### T-Cell Chemotaxis and Infiltration Assay

Mimetas 3-lane 400 um Organoplate® (MIMETAS) was used for these studies. To seed the plates with collagen (Day −2, indexed to T-cell addition day), 50 uL of DPBS was added into the observation port to facilitate making chip filling visible. To form the extracellular matrix barrier, Rat tail collagen-1 (Cultrex) was mixed with HEPES and 37 g/L NaHCO3 in a 8:1:1 ratio to form a 4 mg/mL collagen-1 solution. These components were mixed well > 20 times, being careful not to generate bubbles. Within 10 minutes, 1.8 uL gel solution was seeded into each chip using an automatic repeater pipette (Sartorius). The Organoplate® was then placed in a humidified incubator (37°C, 5% CO2) for 15 minutes to allow polymerization of the collagen-1 gel. 30 uL PBS was then added into the gel inlet to hydrate the ECM layer prior to returning the plate to the incubator. To form the 3D endothelium, HUVECs were trypsinized, resuspended in endothelial medium, counted using an automated cell counter (ViCell Blu, Beckman Coulter), and resuspended to a cell seeding density of 10e6 cells/mL. PBS was removed from the gel inlets, and 2 uL of cell suspension was deposited into the top inlet port using the automatic repeater pipette. Cell suspension was regularly mixed in order to ensure homogenous cell seeding density. After, 50 uL of endothelial medium was added to the same top medium inlet in which the cells were deposited. The Organoplate® was placed with the lid forming a 75 degree angle against the plate stand, and left in this orientation for around 3 hours in order to allow cells to attach. After cell attachment, 50 uL of endothelial medium was added into the top medium outlet. The plate was then placed on the OrganoFlow®, set to an inclination of 7° and an interval of 8 minutes, in a humidified incubator.

On Day −1, tumor cells or empty medium were seeded into the bottom channel using a different seeding strategy. Tumor cells (HCC0827) were trypsinized, resuspended in endothelial medium, counted, and resuspended to a cell seeding density of 10e6 cells/mL. 2 uL of cell suspension was then deposited into the bottom inlet port using the automatic repeater pipette. Cell suspension was regularly mixed in order to ensure homogenous cell seeding density. The Organoplate® was placed with the lid forming a 75 degree angle against the plate stand, but here with the plate rotated 180 degrees from the previous HUVEC seeding step (i.e. top of the plate on the bottom, touching the incubator shelf), and left in this orientation for around 3 hours in order to allow cells to attach. After, 50 uL of endothelial medium was added into the inlet of the bottom perfusion channel, and placed back on the OrganoFlow® rocker.

On Day 0, T-cells or empty medium controls were seeded into the OrganoPlate®. T-cells were harvested gently, centrifuged at 300 x g for 5 minutes, counted, and incubated in dye solution, either 2.5 uM CellTracker Orange CMRA (ThermoFisher) or 1:1000 NucLight Rapid Red (Sartorius), in AIM V medium. For Nuclight Rapid Red dyed cells, cells were dyed at a concentration of 1e6 cells/mL, with no more than 3e6 cells per falcon tube. Conicals of cells in dye solutions were wrapped in foil and placed in an incubator for 30 mins. Halfway through the incubation period, the tubes were inverted several times to gently mix. T-cells were then centrifuged and pelleted to wash out the stain, and resuspended in Complete Assay Medium containing AIM V Medium, 20 IU/mL, 5 ng/mL VEGF and 5 ng/mL bFGF. Cells were then counted and diluted to desired concentration in Complete Assay Medium in order to deliver the number of T-cells per chip indicated in these studies in 50 uL of medium. At this stage, the top medium inlets and outlets were aspirated. 50 uL of T-cell solution was added into the top medium inlet, and 50 uL Complete Assay Medium was added into the top medium outlet. Then, the bottom medium inlet and outlets were aspirated, and replaced with 50 uL medium each containing specified chemokine trigger or control medium solutions. For studies corresponding to Fig 3-5, a half-volume medium re-addition was implemented, in which 25 uL of additional Complete Assay medium were added into the top channel inlet and outlet, and 25 uL of chemokine trigger solution were added into the bottom channel inlet and outlet. For antibody blocking experiments, vehicle alone (PBS), IgG_1_ antibody control (30 ug/mL, R&D Systems, MAB002), ICAM-1/CD54 (10ug/mL, R&D Systems, BBA3) blocking antibody, or VCAM-1/CD106 (30ug/mL, R&D Systems, BBA5) blocking antibody was added into the top channel inlets and outlets at the same time as chemotactic trigger addition into the bottom compartment (Day 0) and also with the half medium refresh (Day 2).

### T-Cell Imaging and Quantification

For data obtained in Figures 1-2 and Supplementary Figures 1-6, images were acquired using a spinning disc confocal and infiltrating T-cells were quantified using a custom FIJI macro as previously described[18].

For data obtained in Figures 3-5 and Supplementary Figures 7-8, imaging was performed either on EVOS microscope or a GE InCell 6500 high content confocal imaging system. Confocal z-stacks acquired were converted into maximum intensity projection images, which were used for analysis. Analyses were performed manually using ImageJ or using a custom python script. For analysing migration distance of T-cells and the number that successfully infiltrate, a python script was developed which utilised the open-source scikit-image library[44]. This analysis pipeline was run in two stages: to accurately identify the PhaseGuidesTM from the brightfield image, and therefore the channel boundaries, and also to identify nuclei that had been stained with DAPI. In order to identify PhaseGuidesTM, a synthetic image that mapped out the position of the PhaseGuidesTM was used as a template to convolve along the image in order to find the position that looked most similar to the distribution of PhaseGuidesTM. To increase the accuracy of this approach, the synthetic image was a 1pixel-width image with intensity bands that are similar to a vertical cross-section of the PhaseGuidesTM (as it is 1 pixel wide, this is less affected by rotation). Fast Normalized Cross Correlation was used for template matching and this led to a processed image with ideally a single horizontal line that had been rotated as per the rotation of the plate. Finding the maximum intensity (and therefore the highest correlation) along the x-axis enabled binarizing the image and then edge detection was used. The original positions of PhaseGuidesTM were then mapped back to this line. Separately, blob detection was used and the distances from the blobs was measured using a signed distance function (i.e. distances are negative if they are behind the line, and positive if they are in front). This meant that channels could be identified just by the sign of the distances. Once the channels had been assigned to each nuclei, it was also possible then to count the number of nuclei in chamber. To assist in detecting the PhaseGuidesTM illumination correction was performed retrospectively by estimating the illumination profile using a low-pass filter (using a Gaussian kernel with a large sigma)[45].

### Barrier Integrity Assays

The barrier integrity of HUVEC endothelial tubes was evaluated before and after the addition of T-cell compatible assay media as previously described[46], and the procedure is detailed within the supplement of this publication[18]. Here, the top chip inlets and outlets were perfused with 0.5 mg/mL 20 kDa FITC Dextran (Sigma, FD20S).

### T-Cell Culture

For activated-only T-cells, CD3+ T-cells were thawed, resuspended in assay medium (RPMI with 10% FBS and 20 IU/mL IL-2), and centrifuged (300xg, 5 mins) (ThermoFisher Scientific; SORVALL ST16, SORVALL LEGEND, or XTR). Cell pellet was resuspended in assay medium and counted using an automated cell counter (Vi-Cell XR, Beckman Coulter). Cells were then diluted to a concentration of 1×10^6 cells/mL in assay medium and then activated using 1:500 TransAct. Cells were then added into a T25 flask and incubated at 37C, 5% CO2 for 72 hours. For activated-rested T-cells, the same procedure was followed as above, except cells were cultured in TransAct for 48 hours. At this time, cells were harvested from flasks, spun down (300xg, 5 mins) and resuspended in assay medium without TransAct. Cells were cultured for an additional 48h rest period.

### Flow Cytometry

Cells were plated at 300,000 cells/well in 96-well U-bottom plates (Corning). Plates were spun down (300xg, 5 mins), washed 1x with 200 uL DPBS (Life Technologies), and spun down again (300xg, 5 mins) to remove supernatant. For live/dead staining, live/dead dye was resuspended as per manufacturer protocol and diluted in PBS at 1:100 dilution. 50 uL of diluted live/dead solution was added to plate wells, mixed thoroughly, and incubated at room temperature for 15 minutes in the dark. Samples were washed 1X with 150 uL PBS and spun down (300xg, 5 mins) to remove supernatant. For Fc blocking and primary antibody staining, 10 uL of Fc block (Miltenyi) were added to each well and incubated for 10 mins in the dark at room temp. Then, 90 uL of antibody cocktail (see details in antibodies and reagents section), prepared in FACs Buffer (Beckton Dickenson) were added to each well and mixed. Samples were incubated for 30 mins at 4C, wrapped in foil to protect from light. Wells were then washed 1X with 100 uL FACS Buffer and 1X with 200 uL FACS Buffer. Plate was then spun down (300xg, 5 mins) and supernatant removed. For sample fixation, 100 uL CytoFIx fixation buffer (Beckton Dickenson) was added to the wells and incubated at 25 mins at room temperature, wrapped in foil to protect from light. Samples were then washed 1X with 100 uL FACS Buffer and 1X with 200 uL FACS Buffer, spun down (300xg, 5 mins) and supernatant removed. Samples were resuspended in 250 uL FACS Buffer and mixed well. Plates were stored at 4C until being read on the cytometer. Staining for compensation controls was conducted on the day of flow analysis as follows. One drop of UltraComp eBeads (eBiosciences) were incubated with 2 uL of the appropriate antibody for 30 mins at room temperature protected from light. For Aqua LIFE/DEAD dye compensation control, 2 drops ArC beads (Life Technologies) were incubated with 2 uL of Live/Dead dye for 30 mins, at room temperature, protected from light. After incubation, beads were washed with flow buffer (500 uL), centrifuged (300xg, 5 mins) and resuspended in 400uL of fresh flow buffer. One drop of ArC negative beads were added to the Aqua tube, and then compensation was run. Flow cytometry was conducted on the LSR Fortessa X-20 (Becton Dickinson), and data was analyzed using FlowJo 10.6.2.

### Immunocytochemistry

Cell cultures in the MIMETAS OrganoPlate® were fixed in 3.7% formaldehyde (Sigma) after 48h, 72h, or 120h in culture and immunostained as previously described[18]. Hoechst 33342 (Thermo Fischer Scientific) was used to stain nuclei. Primary and secondary antibodies were used to stain fixed cultures using products detailed in the antibodies and reagents section.

### Statistical Analysis

Statistical analyses were performed using GraphPad Prism version 8.1.2 (332) for Windows, GraphPad Software, San Diego, California USA, www.graphpad.com. Data were tested for homogeneity in standard deviations, and were square root transformed if needed. Statistically significant differences between means of two or more groups were evaluated using one-way ANOVA (equal variance) or Brown-Forsythe and Welch ANOVA (Gaussian, unequal variance), with multiple comparisons corrected using Dunnett’s, Tukey’s, or Sidak’s. Differences were considered significant if p < 0.05. s

