## Supplementary Information for "Lung Tumor Microphysiological System with 3D Endothelium to Evaluate Modulators of T-Cell Infiltration"

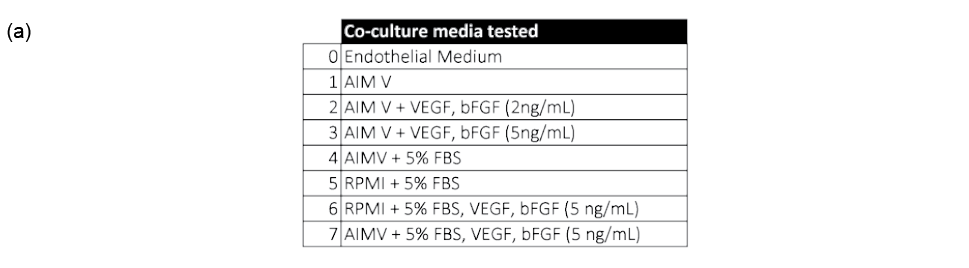

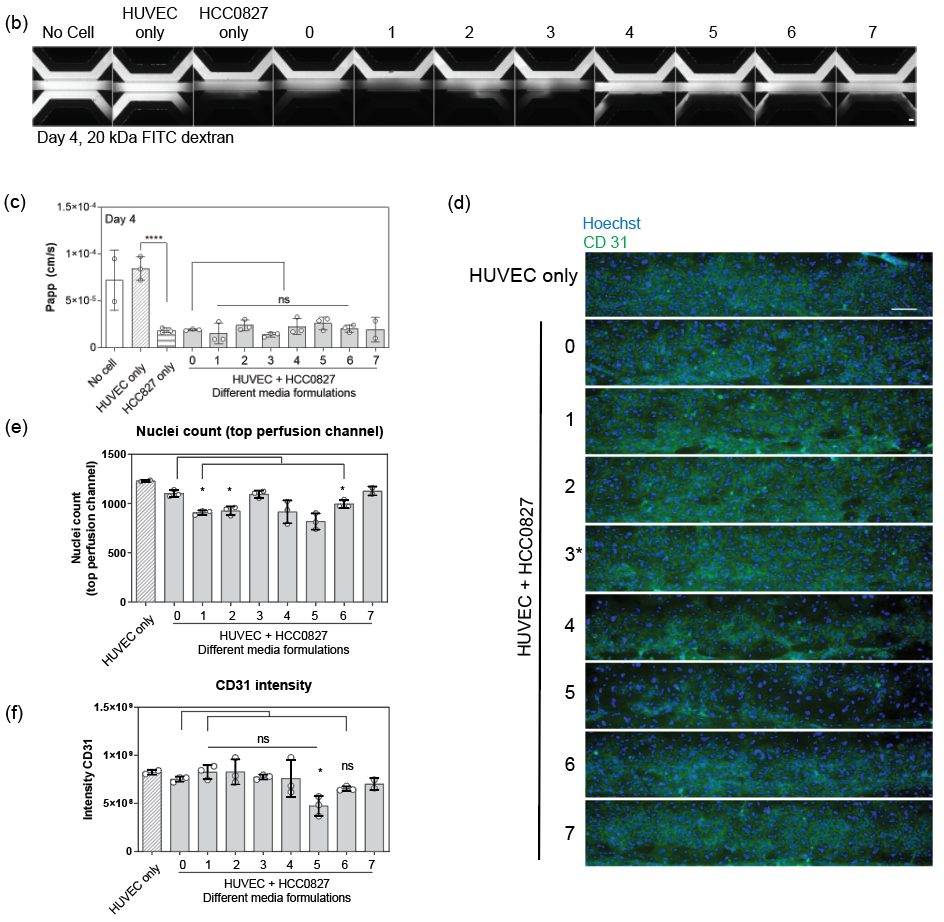

**Supplementary Figure 1: Triculture assay medium optimization studies.** (a), Different assay medium formulations evaluated and compared against endothelial medium. As depicted in Fig 1, chips were cultured in endothelial medium for days -2 to 0, were switched to the indicated medium on day 0, and were cultured until a day 2 endpoint. (b), Example day 2 images of 20 kDa FITC dextran diffusion, introduced immediately prior to imaging, of the conditions as indicated. Included here are no cell, endothelial barrier only, and tumor barrier only controls. Scale bar is 200 μm. (c), Barrier permeability coefficient measurements on Day 2, calculated from images such as so those shown in (b). (d), Representative images of Hoechst and CD31 staining on 3D endothelium in the culture conditions indicated, where scale bar is 100 μm. For images in (d), (e) and (f) depict image-based quantifications of nuclei count and CD31 expression. For (c), (e), and (f), significant differences between medium conditions and endothelial medium controls are shown, with *p< 0.05, ** p<0.01, **** p<0.0001. In (c), Square root transformed data used with ANOVA corrected for multiple comparisons. In (e) and (f), Brown-Forsythe and Welch ANOVA corrected for multiple comparisons and assuming unequal SDs utilized for analysis). N = 3 replicates for all conditions used in statistical testing and HUVEC only controls. N = 2 for no cell control and medium formulation 7, which were not included in statistical testing.

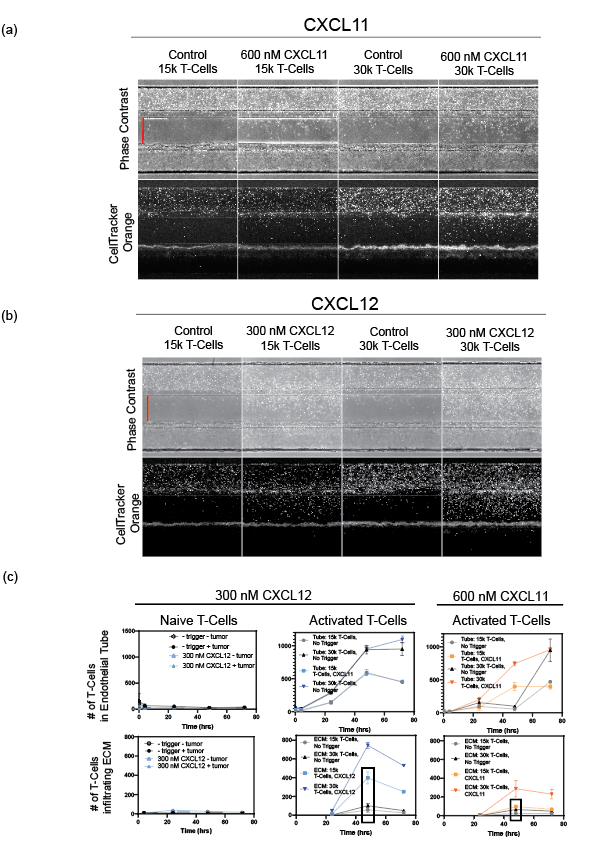

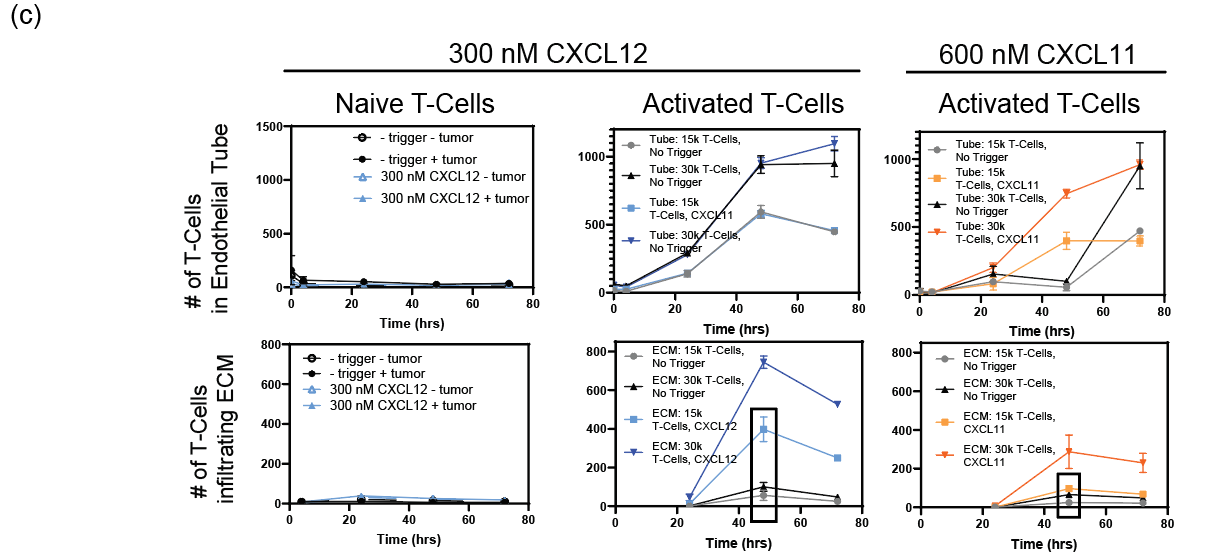

**Supplementary Figure 2: T-cell infiltration by activation status, seeding density, and chemokine type.** (a) and (b), Representative phase contrast and fluorescent imaging at 48h depicting activated T-cell responses in the indicated conditions. (c) Quantified summary of number of T-cells adhering to the endothelial tube and infiltrating the matrix compartment over time, by activation status, seeding density, and chemokine type. Markers indicate means and bars indicate SDs for n = 3 chips. In (a) and (b), middle channel width is 350 μm as indicated by red bars.

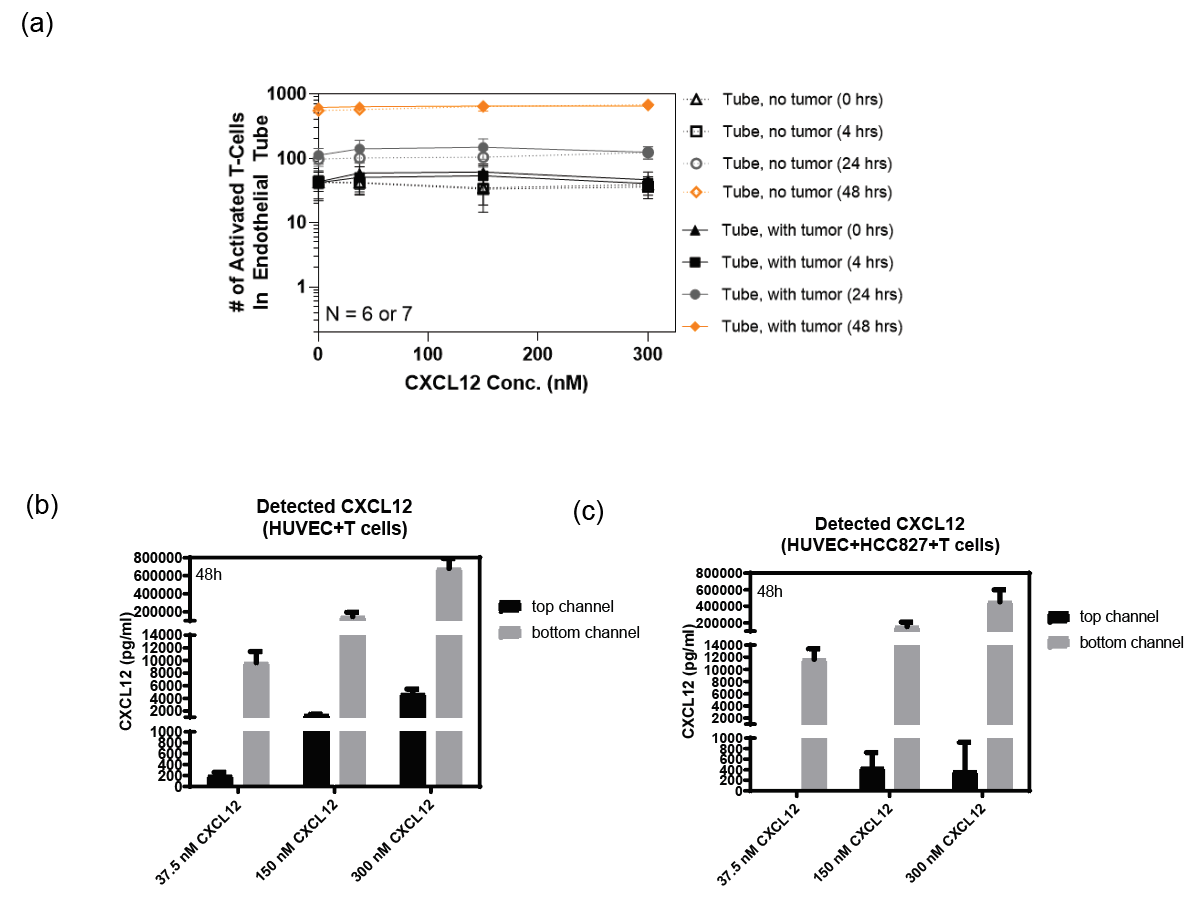

**Supplementary Figure 3: Tumor on chip with 3D endothelium dose-response characterization to CXCL12 chemokine.** (a) T-cell adherence to endothelial tube by CXCL12 dose, shown also by time point and tumor barrier presence/absence. (b) and (c), ELISA data depicting concentration of chemokine in the top endothelial channel and bottom tumor compartment 48 hours after the indicated dose was added into the bottom tumor compartment. Data in (b) were obtained from the no-tumor barrier version of the assay, whereas data in (c) were obtained in using the with tumor barrier configuration. In (b) and (c), bars indicate means and error bars indicate SDs for 4 replicates each.

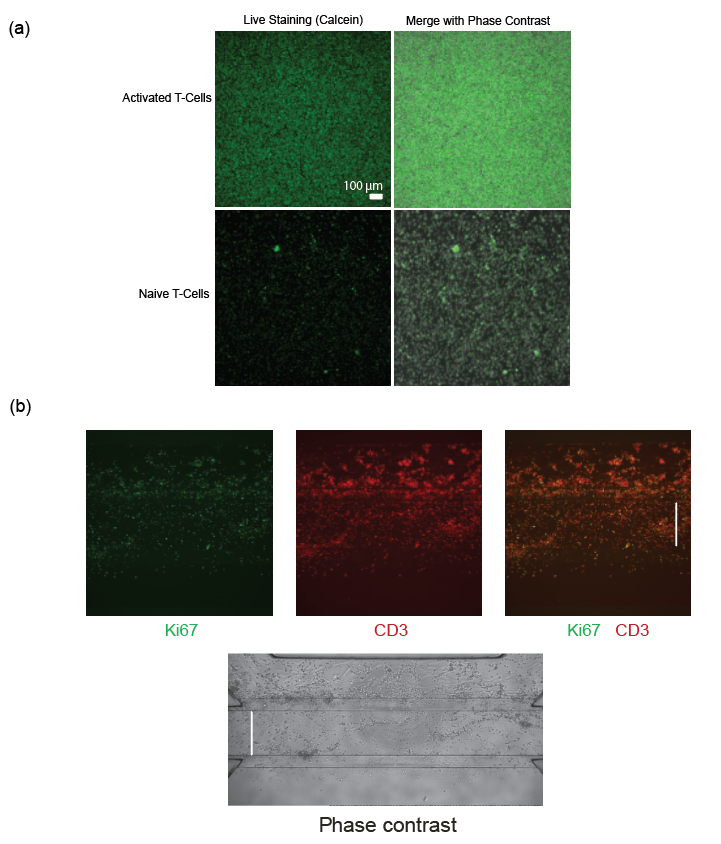

**Supplementary Figure 4: T-cell proliferative burst following activation, both off-chip and in-chip.** (a), Live Calcein staining and phase contrast images of both naïve and activated T-cells, cultured off-chip for 72 hours after activation, with scale bar as indicated. (b) Ki67 (green), proliferation marker, and CD3 (red), T-cell marker, staining of the microphysiological tumor chips with activated T-cells after 72 hours of incubation, along with a phase contrast image depicting the same. Middle channel width is 350 μm as indicated by the vertical bars.

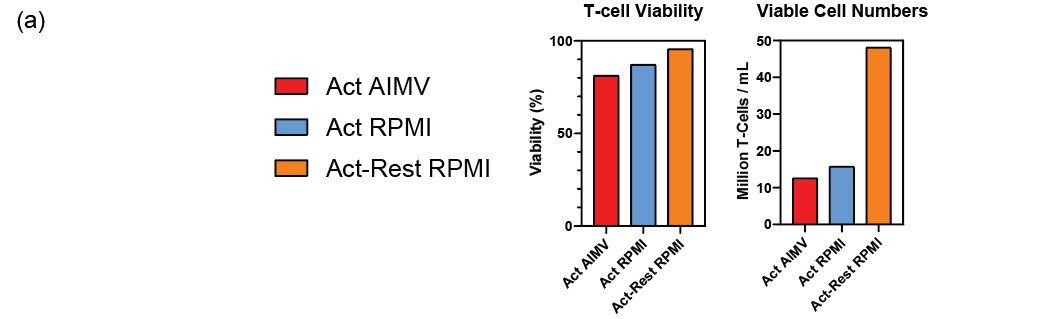

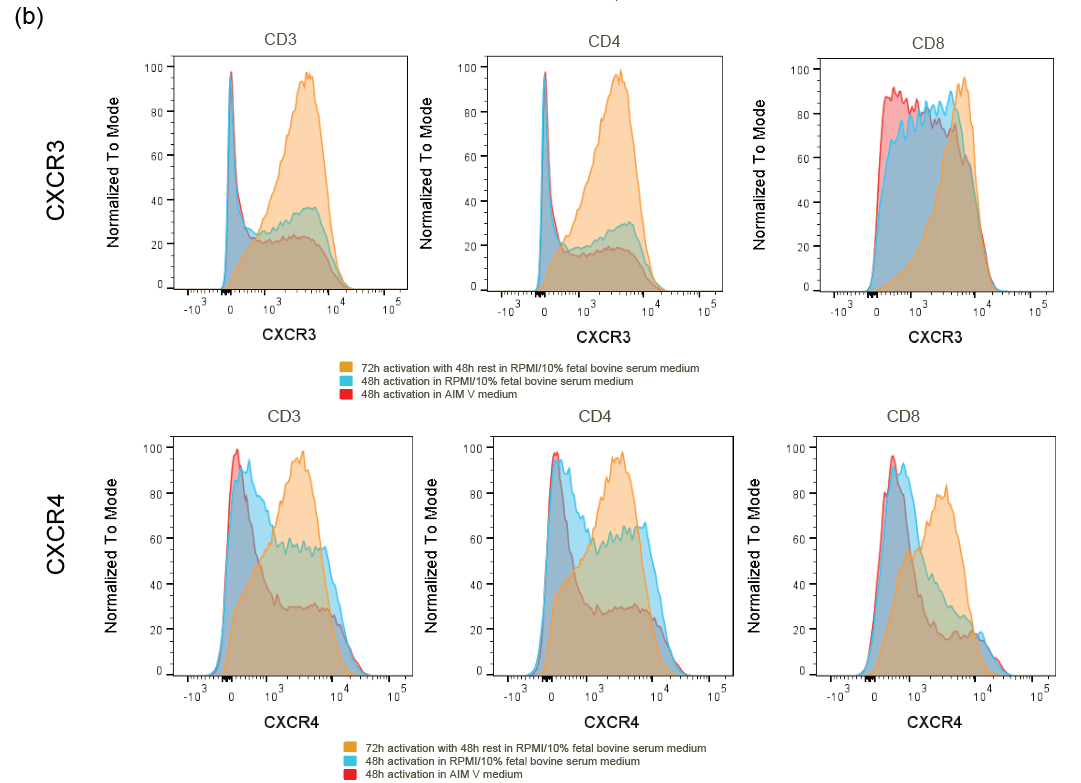

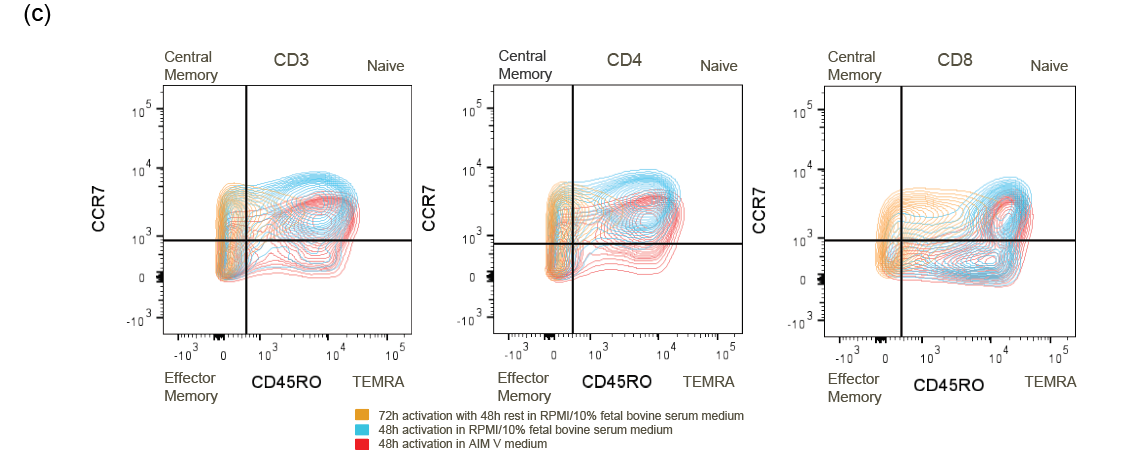

**­­**

**Supplementary Figure 5: T-cell concentrations, receptor expression, and phenotype by T-cell activation protocol.** (a) Comparison of viability and viable cell counts T-cells cultured using activated-only (in AIM V or RPMI-based medium, red and blue respectively) or activated-rested protocol (in RPMI-based medium, orange). Data depict average values obtained over 100 individual images using an automatic cell counter. (b) and (c), Flow cytometry data showing CXCR3 and CXCR4 receptor expression and phenotype by T-cell subset for the same 3 T-cell activation conditions.

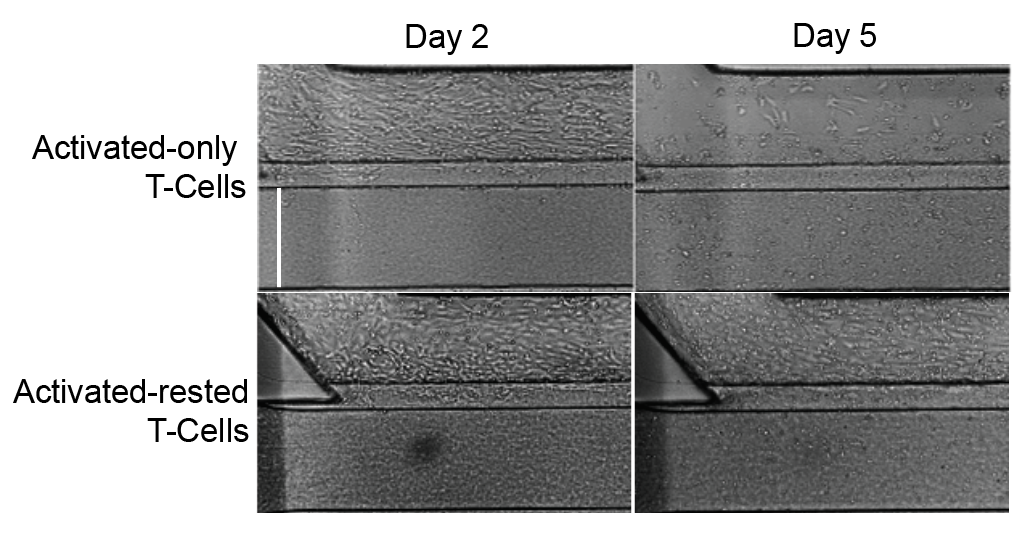

**Supplementary Figure 6: 3D endothelial presence by day and T-cell activation protocol.** Representative brightfield images. Middle channel width is 350 μm as indicated by the vertical bar.

Table 1: Characterization of T-cell populations, prepared using 3 different protocols, using flow cytometry

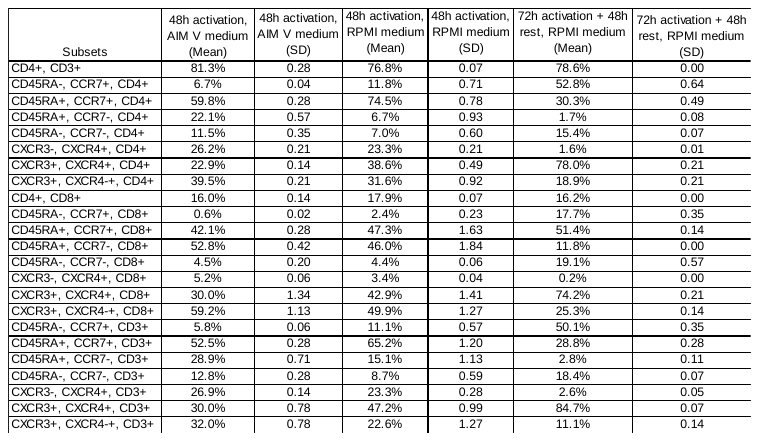

*Equipment, Consumables, Reagents and Antibodies*

**Table 2**: Equipment and consumables.

| Small Equipment and Consumables | Supplier | Details | Lot Number / Other Notes |
| --- | --- | --- | --- |
| OrganoFlow® S or L (MIMETAS, MI-OFPR-S or MI-OFPR-L) | Fisher Scientific | NC1878412, NC1741958 |  |
| Sartorius Biohit PICUS Single Channel Electronic Pipet | Fisher Scientific | Fisher Cat# 14-559-350 Sartorius 735021 |  |
| Corning Costar Aspirator | Fisher Scientific  Corning | 07-200-564  4930 |  |
| Corning Costar Adapter: Multichannel Aspirator | Fisher Scientific  Corning | 07-200-710  4931. |  |
| Zerostat antistatic treatment de-ionizing gun | Sigma | Z108812 | Helpful to use one of these in the winter months, when the lab is dry and full of static, as dispensation of small volumes of hydrogel will not be effective. |
| 3-lane 400μm OrganoPlate™ | MIMETAS | 4004-400-B |  |
| MicroClime Environmental Lid Sterile | Labcyte | LLS-0310 | Recommended to reduce plate evaporation. |

**Table 3**: Biological materials.

| Biological Material | Supplier | Details | Lot Number / Other Notes |
| --- | --- | --- | --- |
| HUVECs | Lonza | Catalog # C2519A. |  |
| Peripheral Blood, Cryopreserved, CD3+ Pan T Cells, Negatively Selected, 25M | ALLCELLS | PB009-1F | We also isolated these in-house using ALLCELLS Leukopaks. |
| HCC0827 | University of Texas Southwestern |  |  |

**Table 4**: Chemical and Molecular Reagents.

| Chemical and Molecular Reagents | Supplier | Details | Lot Number / Other Notes |
| --- | --- | --- | --- |
| Sterile 1X D-PBS (w/o Ca2+, Mg2+) | Life Technologies | 14190-136 |  |
| NaHCO3 | Sigma | S5761-500G |  |
| 1M Tris, pH 7.6 | Bioland Scientific LLC (Fisher Scientific) | NC1546267 |  |
| Collagen-1, 5 mg/mL | R&D Systems | AMSbio Cultrex 3D collagen I rat tail, #3447-020-01. |  |
| 1M HEPES, pH 7.2-7.5 | ThermoFisher Scientific | 15630-080 |  |
| CXCL12 | R&D Systems OR Peprotech | 350-NS-050 (R&D);  300-28A (Peprotech) |  |
| CXCL11/I-TAC | R&D Systems OR Peprotech | 672-IT-025 (R&D);  300-46 (Peprotech) |  |
| CXCL12 Duoset ELISA | R&D Systems | DY350-05, DY008, DY005 |  |
| IL-2 | Miltenyi | 130-097-746 | Use specific activity of each lot and dilute stock to 100 IU/mL |
| BSA | Sigma Aldrich | 10711454001 |  |
| RPMI | Gibco | A42755SA or A10491-01 |  |
| AIM V Medium | Gibco, ThermoFisher Scientific | 12055091 |  |
| COMPLETE HUMAN ENDOTHELIAL CELL MEDIUM /W KIT – 500 ML | Cell Biologics | H1168 |  |
| VEGF | VWR  PeproTech | 10771-972  100-20 |  |
| bFGF | PeproTech | 100-18B |  |
| FBS | Gibco | 1099-141 |  |
| TransAct | Miltenyi | 130-111-160 | Used at 1:500. |
| Nuclight Rapid Red Dye | Essen Bioscience (Manufacturer); Fisher Scientific | 4717 (Manufacturer); Fisher Scientific NC1404054 | Used at 1:1000. |
| CellTracker Orange CMRA Dye | ThermoFisher Scientific | #C34551 |  |
| CellTracker Green | ThermoFisher Scientific | #C2925 |  |
| FITC-dextran 20 kDa | Sigma | FD20S | Stock solution 25 mg/mL in HBSS. |
| Formaldehyde | Sigma | 252549 |  |
| Hoechst 33342 | ThermoFisher Scientific | H3570 |  |
| Anti-CD31 | Dako/Agilent | M0823, Clone JC70A | 1:50 |
| Anti-CD3 | Abcam | ab5690 | 1:100 |
| Ki67 | Cell Signaling Technology | 9449S, clone 8D5 | 1:800, O/N incubation |
| Mouse IgG1, Clone #11711R | R&D Systems | MAB002 | 30 ug/mL |
| VCAM-1/CD106 | R&D Systems | BBA5 | 30 ug/mL |
| ICAM-1/CD54 | R&D Systems | BBA3 | 10 ug/mL |
| Goat-anti-mouse Alexa Fluor PLUS 488 | ThermoFisher | A32731 |  |
| Goat-anti-mouse Alexa Fluor PLUS 555 | ThermoFisher | A32727 |  |
| Goat-anti-mouse IgG (H+L) Alexa Fluor 647 | ThermoFisher | A21236 |  |
| Goat-anti-rabbit Alexa Fluor PLUS 488 | ThermoFisher | A32731 |  |
| Goat anti-Rabbit IgG (H+L) Alexa Fluor Plus 555 | ThermoFisher | A32732 |  |
| Donkey Anti-Rabbit IgG Polyclonal Antibody (CF™ 647) | Bio-Connect | 20047 |  |

**Table 5**: T-cell isolation reagents

| Reagents | Supplier | Details | Lot Number / Other Notes |
| --- | --- | --- | --- |
| Custom PBMC Isolation Kit | Miltenyi | 130-123-456 |  |
| Pan-T Isolation Kit | Miltenyi | 130-096-535 |  |
| CS 10 | BioLife Solutions | 210102 |  |

**Table 6**: Flow Cytometry Reagents.

| Reagents | Supplier | Details | Lot Number / Other Notes |
| --- | --- | --- | --- |
| DPBS | Life Technologies | Cat. 14190-136 |  |
| FcR Blocking Reagent | Miltenyi | Cat. 130-059-901 | Lot 5200809006 |
| BD Pharminen Stain Buffer (BSA) | Beckton Dickenson | Cat.554657 | Lot 9263284 |
| BD CytoFix | Beckton Dickenson | Cat. 554655 | Lot 9179165 |
| UltraComp eBeads | eBiosciences | #01-2222-42 |  |
| ArC amine-reactive compensation bead kit | Life Technologies | #A10346 |  |

**Table 7**: Flow Cytometry Antibodies

| Marker | Color | Clone | Host | Isotype | Vendor | Cat# (1:100 dilution) | Isotype  Cat# |
| --- | --- | --- | --- | --- | --- | --- | --- |
| Live Dead/Aqua | BV510 |  |  |  | Invitrogen | L34966 |  |
| CXCR3 | BV421 | G025H7 | Mouse | IgG1, κ | BioLegend | 353716 | 400158 |
| CXCR4 | APC | 12G5 | Mouse | IgG2a, κ | BioLegend | 306510 | 400220 |
| CD3 | AF488 | UCHT1 | Mouse | IgG1, K | BioLegend | 300415 | 400129 |
| CD4 | PE-Cy7 | OKT4 | Mouse | IgG2b, κ | Biolegend | 317414 | 400326 |
| CD8 | AF700 | SK1 | Mouse | IgG1, κ | BioLegend | 344724 | 400144 |
| CCR7 | Percp-Cy5.5 | G043H7 | Mouse | IgG2a, κ | BioLegend | 353220 | 400252 |
| CD45RA | BV605 | HI100 | Mouse | IgG2b, κ | BioLegend | 304134 | 400350 |
